# Programmable Microbial Ink for 3D Printing of Living Materials Produced from Genetically Engineered Protein Nanofibers

**DOI:** 10.1101/2021.04.19.440538

**Authors:** Anna M. Duraj-Thatte, Avinash Manjula-Basavanna, Jarod Rutledge, Jing Xia, Shabir Hassan, Arjirios Sourlis, Andrés G. Rubio, Ami Lesha, Michael Zenkl, Anton Kan, David A. Weitz, Yu Shrike Zhang, Neel S. Joshi

## Abstract

Living cells have the capability to synthesize molecular components and precisely assemble them from the nanoscale to build macroscopic living functional architectures under ambient conditions.^1–3^ The emerging field of living materials has leveraged microbial engineering to produce materials for various applications, but building 3D structures in arbitrary patterns and shapes has been a major challenge.^1–14^ We set out to develop a new bioink, termed as “microbial ink” that is produced entirely from genetically engineered microbial cells, programmed to perform a bottom-up, hierarchical self-assembly of protein monomers into nanofibers, and further into nanofiber networks that comprise extrudable hydrogels. We further demonstrate the 3D printing of functional living materials by embedding programmed *Escherichia coli* (*E. coli*) cells and nanofibers into microbial ink, which can sequester toxic moieties, release biologics and regulate its own cell growth through the chemical induction of rationally designed genetic circuits. This report showcases the advanced capabilities of nanobiotechnology and living materials technology to 3D-print functional living architectures.

---

3D bioprinting technology, which is relatively well-established for printing mammalian cells in the context of tissue engineering, has more recently been applied to print microbial cells for biotechnological and biomedical applications.^9–16^ Although inkjet printing, contact printing, screen printing, and lithographic techniques have been explored to print microbes, extrusion-based bioprinting has become one of the most widely used techniques due to its simplicity, compatibility with a variety of bioinks and cost-effective instrumentation.^16–19^ In an early example of this concept, a mixture of alginate and *E. coli* was extruded onto a printing surface consisting of calcium chloride, upon which the alginate molecules crosslink to form a solidified gel.^14^ A similar ionic crosslinking strategy was exploited to generate photocurrent with 3D printed cyanobacteria.^20^ In another approach, a multi-material bioink comprised of hyaluronic acid, *κ*-carrageenan, fumed silica, and a photo-initiator was employed to 3D-print *Pseudomonas putida* and *Acetobacter xylinum*. Also, photo-crosslinked pluronic F127 acrylate-based bioinks have been utilized to print living, responsive materials/devices and catalytically active living materials.^11,12,21^

An alternative strategy made use of freeze-dried *Saccharomyces cerevisiae* as the primary component of a bioink formulation consisting of nanocellulose, polyethylene glycol dimethacrylate, and a photoinitiator.^13^ The latter approach yielded remarkably high cell densities of 10^9^ cells ml^−1^, but the need for freeze-drying could significantly affect the survival rate of other microbial species as well as their thixotropic behavior. In an interesting approach, the viscoelastic gel-like characteristics of *Bacillus subtilis* (*B. subtilis*) biofilms facilitated direct printing. However, the wild-type biofilms were unable to maintain the printed line widths (as they expanded three-fold in width after printing), while the engineered variants had lower storage modulus and viscosity that restricted their printing in multiple layers.^10^ In yet another strategy, a fused deposition modeling was adapted to deposit molten agarose (75 °C) containing *B. subtilis* spores onto a cold substrate (16 °C), resulting in hardened patterns upon cooling.^9^ Here, the high-temperature processing works well for spores, but limits applicability to a wide range of cell types.

Although the above examples demonstrate that many bioink designs have already been explored, none so far have fully leveraged the genetic programmability of microbes to rationally control the mechanical properties of the bioink. This would be advantageous for several reasons, including the possibilities of more sustainable manufacturing practices, raw material fabrication in resource-poor environments (terrestrial or extra-terrestrial), and enhanced material performance through bio-inspired design and the precision of genetic programming. In contrast to the examples described above, we envisioned to 1) design a new extrudable bioink that had high print fidelity, 2) produce the bioink entirely from engineered microbes by a bottom-up approach and 3) create a programmable platform that would enable novel functions for the macroscopic 3D living materials.

There is a critical need to develop advanced bioinks with tunable mechanical strength, high cell viability, and high print fidelity.^19^ A printable bioink requires a viscosity low enough to facilitate extrusion, but high enough to retain its shape after printing.^16^ In this regard, shear-thinning hydrogels, which decrease their viscosity with increasing shear stress, are an attractive option. Moreover, it should be noted that bioinks are biocompatible materials typically meant to recapitulate an extracellular matrix (ECM) to provide a congenial environment for the growth of living cells with predefined structures and functions. We envisioned that instead of embedding microbes in an ECM-mimicking bioink, we could repurpose the ECM of the microbial biofilm itself to serve as a programmable bioink. This idea is built on our earlier work, wherein we showed that the native proteinaceous curli nanofibers of an *E. coli* biofilm ECM can be genetically engineered by fusing functional peptides/proteins to the curli CsgA monomer to produce a shear-thinning hydrogel.^4^ However, in order to create a bioink with the desired viscoelastic performance, we introduced a genetically-programmed crosslinking strategy, inspired by fibrin (**Fig. 1**).^22^ Fibrin is a protein involved in the clotting cascade, which activates its polymerization to form blood clots. Fibrin’s polymerization is driven in part by noncovalent interactions between an alpha-chain domain present on the N-terminus of one fibrin monomer (i.e. the “knob” domain) and a gamma-chain domain on the C-terminus (i.e. the “hole” domain) of an adjacent monomer.^22^ Our microbial ink design repurposes this binding interaction between alpha and gamma modules, *i.e.* the knob-hole interaction, to introduce non-covalent crosslinks between nanofibers and enhance mechanical robustness while maintaining shear-thinning properties (**Fig. 1**).

**Figure 1.**
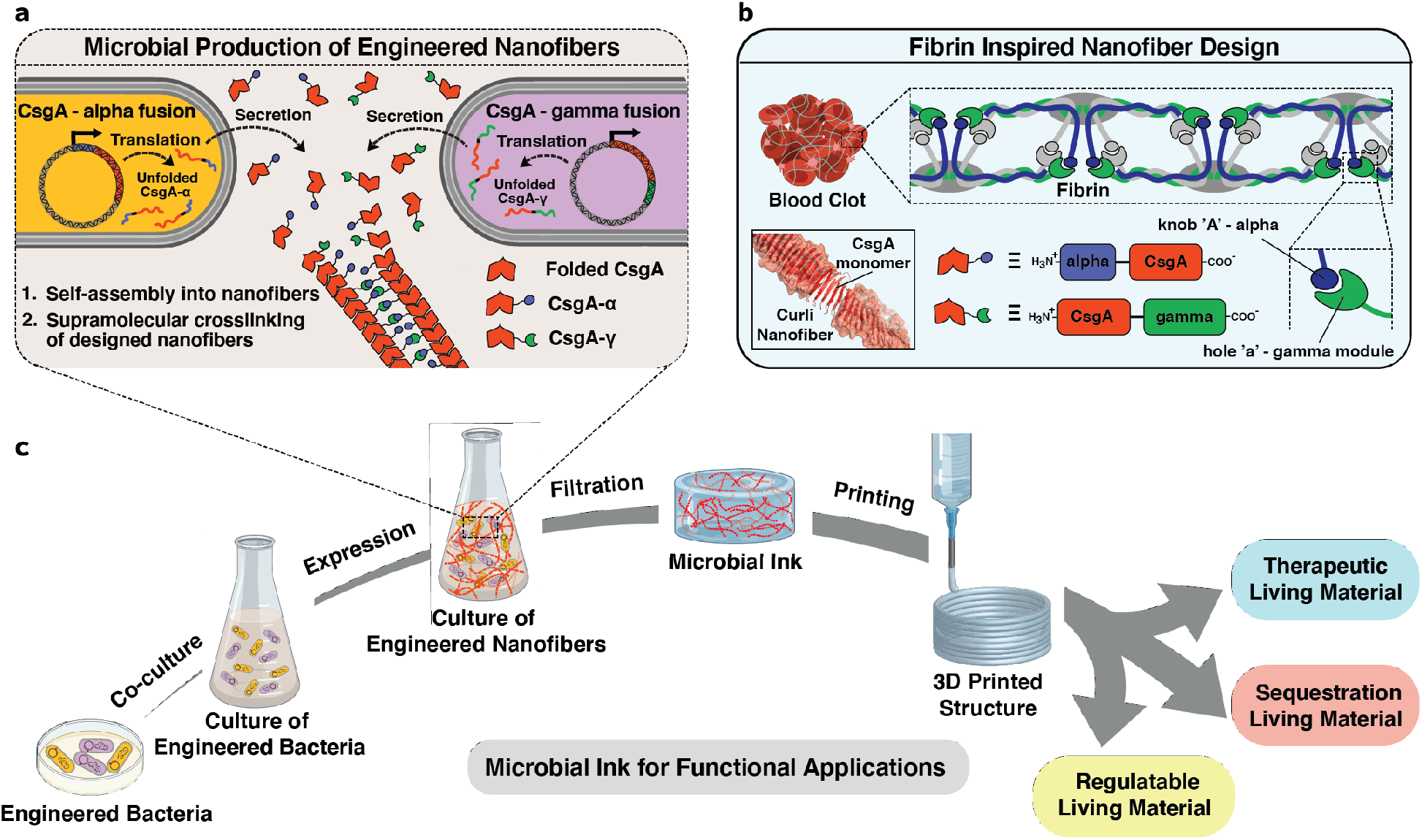
Schematics of the design strategy, production, and functional applications of microbial ink. **a.** *E. coli* was genetically engineered to produce microbial ink by fusing *α* (knob) and *γ* (hole) protein domains, derived from fibrin, to the main structural component of curli nanofibers, CsgA. Upon secretion, the CsgA-*α* and CsgA-*γ* monomers self-assemble into nanofibers crosslinked by the knob-hole binding interaction. **b.** The knob and hole domains are derived from fibrin, where they play a key role in supramolecular polymerization during blood clot formation. **c**. The protocol for the production of microbial ink from the engineered protein nanofibers involves standard bacterial culture, limited processing steps, and no addition of exogenous polymers. Microbial ink was 3D printed to obtain functional living materials.

Using the Biofilm Integrated Nanofiber Display (BIND) technology developed in our laboratory,^23^ we genetically grafted the “knob” and “hole” protein domains to the N- and C-terminus of CsgA, respectively, to create the fusion proteins CsgA-*α* and CsgA-*γ* (**Fig. 1, Supplementary Table 1**). CsgA-*α* and CsgA-*γ* were expressed separately in engineered *E. coli* strain PQN4 along with the auxiliary curli genes necessary for secretion and assembly, and the resulting curli nanofibers were imaged using transmission electron microscopy (TEM). After staining with 1% uranyl formate, the nanofibers showed diameters of ~5.5 nm (CsgA-*α*) and ~6.7 nm (CsgA-*γ*) (**Fig. 2a**). Notably, curli nanofibers composed of wild-type CsgA have diameters of ~4 nm.^23^ Thus, the observed trend in nanofiber diameters is qualitatively consistent with the relative sizes of the fused domains – 11 amino acids for “knob” (CsgA-*α*) and 127 amino acids for “hole” (CsgA-*γ*). When the two types of *E. coli* cells, each expressing either CsgA-*α* or CsgA-*γ*, were co-cultured (CsgA-*αγ*), they produced nanofibers that display “knob” and “hole” domains. TEM imaging showed three nanofiber populations with diameters of ~5.5 nm, ~6.7 nm, and ~10 nm (**Fig. 2a, Supplementary Figure 1**). We attribute the 10 nm diameter nanofibers to supramolecular crosslinking mediated by noncovalent interactions of the “knob” and “hole” domains (**Fig. 1**). We then created hydrogels from the microbial cultures using a simple filtration protocol, as described in our earlier reports.^4^ Briefly, the microbial culture was filtered through a nylon membrane to concentrate the curli nanofibers, and then treated with guanidinium chloride, nuclease, and sodium dodecyl sulfate to obtain cell-free hydrogels composed of the designed curli nanofibers (**Fig. 2b**).^4^ Field-emission scanning electron microscopy (FESEM) indicated a fibrous microstructure for all three hydrogels (CsgA-*α*, CsgA-*γ* and the co-culture CsgA-*αγ*), with the fiber alignment suggesting hierarchical assembly through the lateral association of functional curli nanofibers (**Fig. 2b**).

**Figure 2.**
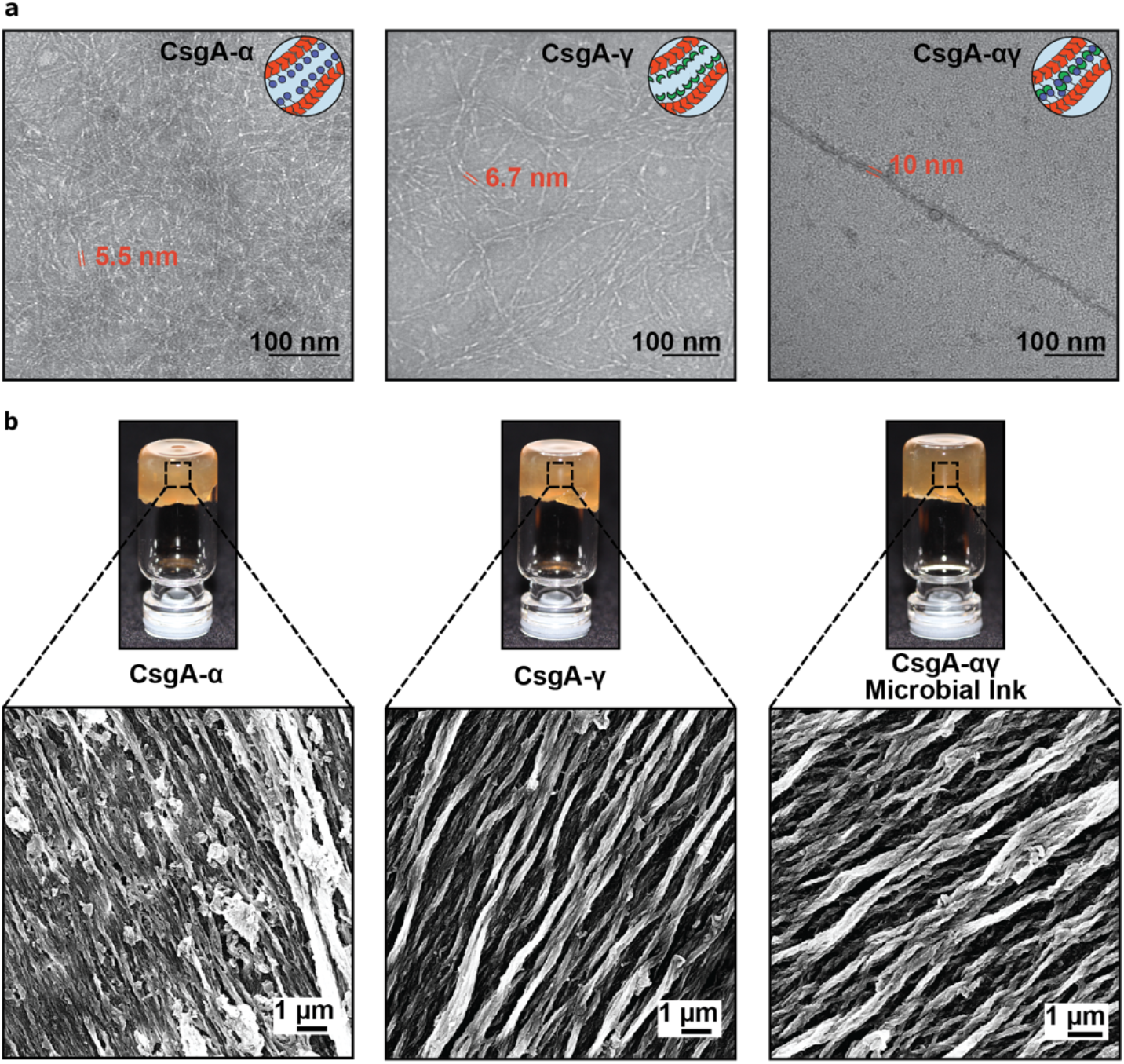
Optical and electron microscopy images of functional curli nanofibers and the corresponding hydrogels. **a**. TEM images of self-assembled nanofibers of CsgA-*α*, CsgA-*γ* and CsgA-*αγ* (co-culture of CsgA-*α* and CsgA-*γ*) after recombinant expression. **b**. Optical images of CsgA-*α*, CsgA-*γ* and microbial ink CsgA-*αγ* hydrogels with the corresponding FESEM images show the presence of aligned microscopic fiber bundles.

We next investigated the rheological properties of CsgA-*α*, CsgA-*γ* and CsgA-*αγ* hydrogels, in order to validate their potential as extrudable bioinks. Frequency sweep experiments revealed that the storage modulus (G’) of the CsgA-*αγ* hydrogel was several-fold higher than that of the CsgA-*α* and CsgA-*γ* hydrogels alone, while the G’ of all the hydrogels were higher than their loss modulus (G”) by an order of magnitude (**Fig. 3a**). Strain sweep experiments showed that the hydrogels were stable up to ~10% strain, above which a crossover point is observed as G’ decreased and G” increased. (**Fig. 3b**). The viscosity of all the hydrogels was also found to decrease with increasing shear rate, which indicates their shear-thinning behavior (**Fig. 3c**). Similarly, the shear modulus (G) of CsgA-*αγ* was higher than that of CsgA-*α* and CsgA-*γ* by 6- and 3-fold, respectively (**Fig. 3d**). The yield stress (*σ_y_*) of CsgA-*αγ* was nearly twice that of CsgA-*α* and CsgA-*γ* (**Fig. 3e**). From all the above experiments, it is clear that supramolecular crosslinking of “knob” and “hole” domains in CsgA-*αγ* significantly increased the G’, G, *σ_y_*, and viscosity, making it better suited than CsgA-*α* or CsgA-*γ* for extrusion printing.^16,19^ On the other hand, when CsgA-*α* and CsgA-*γ* fibers expressed in separate cultures were mixed (CsgA-*αγ*-mix) in a 1:1 volume ratio also yielded hydrogels with rheological properties similar to CsgA-*αγ*, which further confirms our hypothesis about supramolecular crosslinking between the complementary fibers (**Supplementary Figures 2-3**).

**Figure 3.**
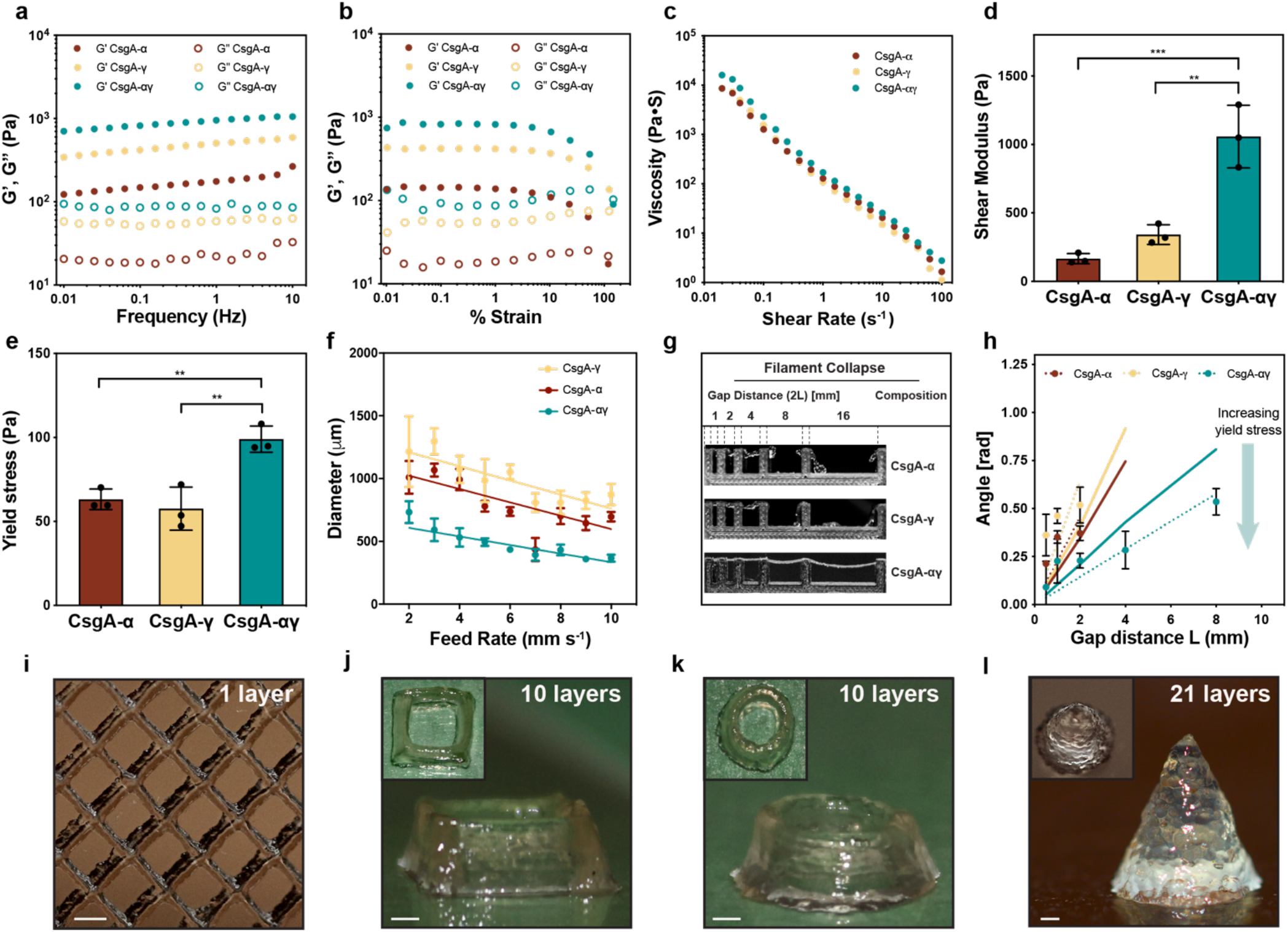
Rheological properties and 3D printing of CsgA-*α*, CsgA-*γ* and microbial ink CsgA-*αγ*. The storage modulus (G’) and loss modulus (G”) under frequency sweep (**a**) and oscillatory sweep (**b**). **c.** Viscosity as a function of shear rate, **d.** Shear modulus and **e.** yield stress. **f.** Printed line diameter as a function of feed rates ranging from 2 to 10 mm s^−1^ at 20 psi pressure. **g.** Images of filament collapse test and **h.** the plot of deflection angle versus pillar gap distances. Experimental data: solid line, theoretically predicted data: dotted line. Data represented as mean ± standard deviation. \*\**p ≤* 0.01, \*\*\**p ≤* 0.001, one-way ANOVA followed by Dunnett’s test. 3D printed structures using the microbial ink CsgA-*αγ* (**i**) single layer grid, (**j**) 10-layer square, (**k**) 10-layer circle, and (**l**) 21-layer solid cone. Insets in **j-l.** are corresponding top views. Scale bar 1 mm.

We then tested the printability of the CsgA-*α*, CsgA-*γ* and CsgA-*αγ* hydrogel-based bioinks using a customized 3D printer (**Supplementary Figure 4**). First, the hydrogel-based bioinks were extruded under a range of feed rates (2-10 mm s^−1^) and pressures (20-40 psi) to understand their printing performance (**Fig. 3f**, **Supplementary Figures 5-10**). The printed line widths of CsgA-*α* and CsgA-*γ* bioinks were nearly two times that of CsgA-*αγ* for the same feed rate and pressure, indicating the superior structural integrity of CsgA-*αγ* (**Fig. 3f**, **Supplementary Figures 5-10**). Subsequently, we tested the shape fidelity of the bioinks according to a published protocol (filament collapse test) to provide a quantitative comparison between bioinks.^24^ A single line (filament) of each bioink was extruded at a nozzle moving speed of 5 mm s^−1^ on a platform with pillars at known gap distances, to bridge the pillar gaps (**Fig. 3g, Supplementary Video 1**). The CsgA-*α* and CsgA-*γ* bioinks were unable to bridge gap distances of 8 mm and above, whereas the CsgA-*αγ* bioink was able to support its own weight for gap distances as large as 16 mm. This remarkable property, along with the higher *σ_y_* and viscosity, suggest that the supramolecular knob-hole crosslinks allow for fast reassembly after extrusion – an important feature for optimal bioink performance.^25,26^ Shape fidelity was assessed quantitatively by measuring the angles of deflection of the overhung bioink fibers under gravitational force (**Fig. 3h**).^24^ When the angles of deflection are plotted against the half gap distances, the resulting slope will decrease for the bioink with higher *σ_y_*.^24^ This experimental data was also consistent with the reported theoretical model, while the deviation of experimental and predicted slopes was also in line with the original report, which observed that the model overestimated the angles of deflection, likely due to the exclusion of gel viscoelasticity and surface tension from the theoretical model.^24^ We then utilized the CsgA-*αγ* bioink to 3D-print defined patterns and shapes. A single-layer grid shows the finer line structures of the printed pattern obtained with a resolution of ~300 μm from a 27G needle (**Fig. 3i**). The multilayered architectures presented in **Fig. 3j** (10-layered square), **Fig. 3k** (10-layered circle) and **Fig. 3l** (21-layered solid cone) demonstrated the structural integrity of the microbial ink.

After demonstrating the printing performance of the microbial ink, we introduced the genetically engineered microbes to the hydrogel to produce 3D-printed living functional architectures. Herein, we present a living material for therapeutic applications, wherein a chemical inducer, isopropyl *β*-D-1-thiogalactopyranoside (IPTG) was utilized to signal the programmed *E. coli* (PQN4-Azu) to synthesize on demand an anticancer biologic drug, azurin, and secrete it into the extracellular milieu (**Fig. 4a, Supplementary Figure 11**).^27^ The microbial ink, laden with PQN4-Azu cells, was used to print a 2D capsule pattern that was incubated in the lysogeny broth (LB) media with (+) and without (−) the inducer IPTG. After 24 and 48 h of incubation, the detection of the secreted azurin by an anti-azurin antibody illustrated the functioning of the 3D-printed therapeutic living material (**Fig. 4a**). Next, we produced a living material designed to sequester a toxic chemical, Bisphenol A (BPA). For this demonstration, we grafted a BPA-binding peptide domain to CsgA (CsgA-BPABP) and loaded the PQN4 cells expressing CsgA-BPABP into the microbial ink (**Fig. 4b, Supplementary Figure 12**).^28^ After printing a 2D pattern with the cell-laden ink, the pattern was incubated in LB media with 1 mM BPA. Liquid chromatography-mass spectroscopy (LC-MS) analysis showed that the microbial ink embedded with CsgA-BPA biofilm sequestered nearly 8% and 27% of the BPA after 12 and 24 h of incubation, respectively, while a negative control pattern made with microbial ink only, showed no appreciable BPA sequestration (**Fig. 4b**). Finally, we show that the cell growth within the printed material can be regulated by inducing a genetic circuit (**Fig. 4c, Supplementary Figure 13**). To accomplish this, *E. coli* (PQN4-MazF) cells were programmed to express (upon induction with IPTG) the endoribonuclease toxin, MazF, that inhibits protein synthesis by cleaving mRNA, and can arrest cell growth and/or lead to cell death.^29^ PQN4-MazF cells in the printed structure were found to proliferate in the absence of IPTG, but after 2 h of IPTG induction, the colony forming unit (CFU) count reduced by nearly two orders of magnitude due to expression of MazF (**Fig. 4c**). However, subsequently, the cell growth was restored to some extent, likely due to the native MazFE toxin-antitoxin system in *E. coli* (**Fig. 4c**).^29^ Such a regulation system can be further engineered to effectively control the cell growth and/or to induce cell death depending on the need.

**Figure 4.**
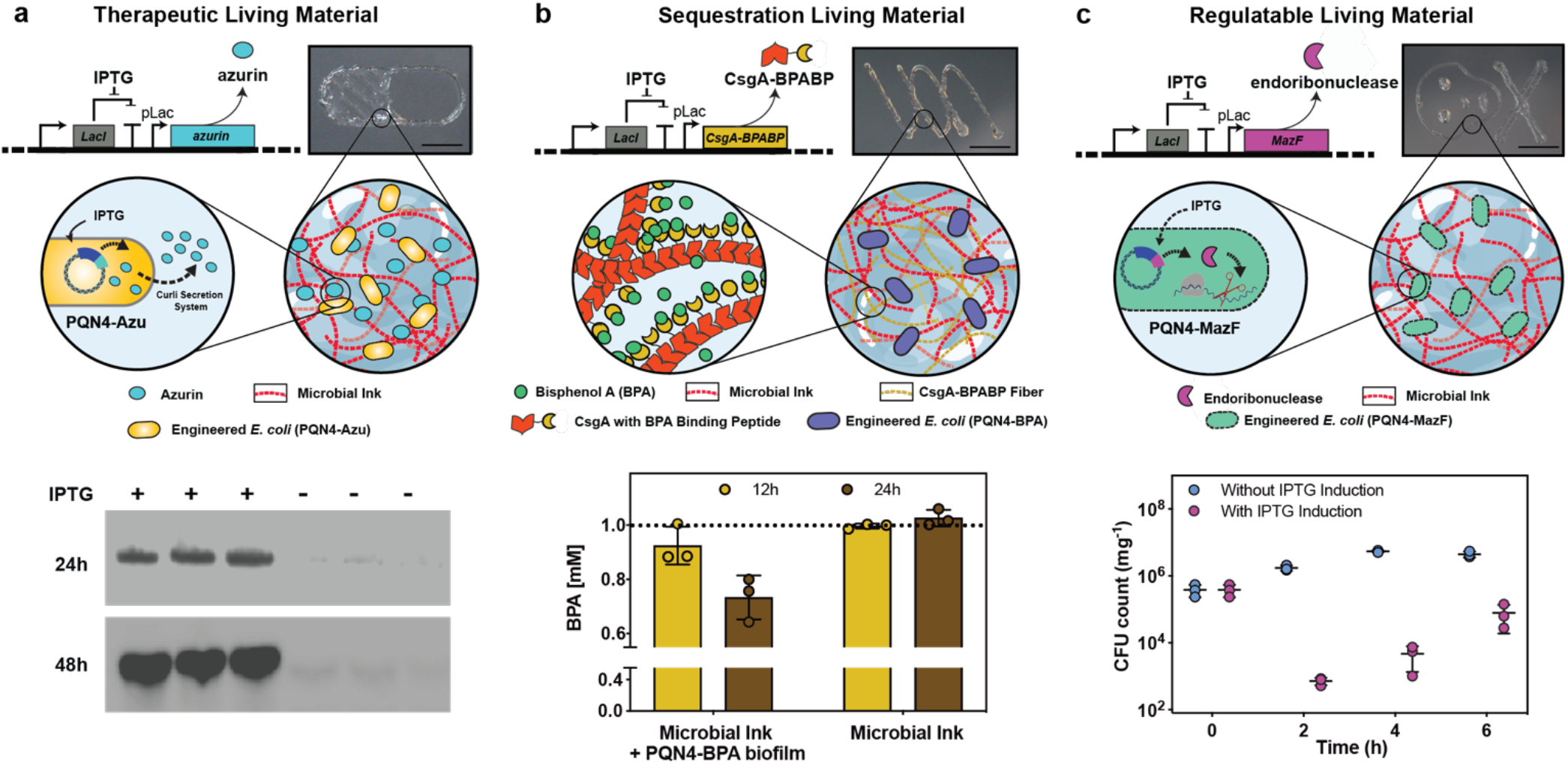
3D printing of functional living materials. **a.** Genetic design of *E. coli* (PQN4-Azu) cells, programmed to secrete an anticancer biologic drug azurin along with image of printed living material (*top*), and the incorporation of PQN4-Azu cells into the CsgA-*αγ* microbial ink (*middle*). Western blot (*bottom*) shows the difference in azurin detected in the supernatant of the printed structure with and without IPTG induction. **b.** Genetic design of *E. coli* (PQN4-BPA), programmed to produce extracellular fibers displaying BPA-binding peptide (CsgA-BPABP) along with image of printed living material (*top*), and incorporation of CsgA-BPABP biofilm into microbial ink (*middle*). After 12 and 24 hours, the BPA concentration (*bottom*) in the supernatant of the printed structure was analyzed by LCMS. Dotted line represents the initial BPA concentration of 1 mM. **c.** Genetic design of *E. coli* (PQN4-MazF) cells, programmed to express an endoribonuclease MazF, that inhibits/arrests cell growth along with image of printed living material (*top*), and incorporation of PQN4-MazF cells into microbial ink (*middle*). CFU count from printed structure over time, with and without IPTG induction. Scale bar 5 mm. Data represented as mean ± standard deviation.

In summary, we have genetically engineered the ECM of *E. coli* biofilms to produce a shear-thinning hydrogel by supramolecular crosslinking of fibrin-inspired recombinant protein nanofibers. Instead of using an external biocompatible material as the bioink, we have shown that a cell-laden bioink with target rheological and functional properties can be created purely through genetic engineering and minimal processing. The printability, structural integrity and print fidelity of the microbial ink was demonstrated with the aid of detailed rheological studies and filament collapse tests. By incorporating programmed microbes/biofilms into the microbial ink, we have 3D-printed living materials that can be chemically induced to release the anticancer drug azurin, remove BPA from their surroundings, and regulate their own cell growth. The microbial ink design can be further customized for various biotechnological and biomedical applications using the ever-growing toolkit of biological parts being developed by synthetic biologists. This microbial manufacturing technology could also be particularly useful for structure building in space or extraterrestrial habitats, where raw material transport is extremely difficult, making on-demand generation of building materials from very limited resources essential.^30^

## Acknowledgements

Work was performed in part at the Center for Nanoscale Systems at Harvard. Work in the N.S.J. laboratory is supported by the National Institutes of Health (1R01DK110770), the National Science Foundation (DMR 2004875) and the Wyss Institute for Biologically Inspired Engineering at Harvard University. Work in the D.A.W. laboratory is supported by the Harvard University Materials Research Science and Engineering Center (NSF Grants DMR-1420570 and DMR-2011754). Work in the Y.S.Z. laboratory was supported by the Lush Prize and the Brigham Research Institute. Parts of the schematics were adapted from BioRender.com.

## Author contributions

A.M.D.-T. and A.M.-B. conceived the project. A.M.D.-T. and J.R. cloned CsgA-*α* and CsgA-*γ*. A.M.D.-T., A.M.-B., A.S. and J.R. produced hydrogel-based microbial inks. A.M.-B. did FESEM and TEM imaging. J.X. tested the rheological properties of the microbial inks. S.H., A.G.R., A.L. and M.Z. tested printing performance and printed 3D structures. D.A.W. supervised J.X. Y.S.Z. supervised S.H., A.G.R., A.L. and M.Z. A.M.D.-T. cloned PQN4-Azu and PQN4-BPA. A.K. cloned PQN4-MazF. A.M.D.-T. and A.M.-B tested and analyzed living materials produced from PQN4-Azu, PQN4-BPA and PQN4-MazF. A.M.-B., A.M.D.-T., and N.S.J. wrote and/or edited the manuscript. All authors discussed and commented on the manuscript.

## Competing interests

A.M.D.-T., A.M.-B., A.S., and N.S.J. are inventors on a patent application submitted by Harvard University.

## Data availability

All relevant data supporting the findings of this study and the plasmids and strains used are available within the Article and its Supplementary Information or from the corresponding authors upon request. Source data are provided with this paper.

