## Supplemental Information for "Programmable Microbial Ink for 3D Printing of Living Materials Produced from Genetically Engineered Protein Nanofibers"

**This file includes:**

Methods
Supplementary Figures 1-13
Supplementary Table 1

### Methods

**Plasmids for production of microbial ink:** The CsgA-$\alpha$ construct was designed by fusing the fibrin derived alpha (knob) peptide to the N-terminus of CsgA with an intervening 12-amino-acid flexible linker. However, in case of CsgA-$\gamma$ construct, the gene encoding the fibrin derived gamma (hole) protein was fused to the C-terminus of CsgA with an intervening 36-amino-acid flexible linker. Both genes encoding CsgA-$\alpha$ and CsgA-$\gamma$ were synthesized (Integrated DNA Technologies) and cloned into pET21d vector using isothermal Gibson assembly (New England Biolabs). Both pET21dCsgA-$\alpha$ and pET21dCsgA-$\gamma$ plasmids were transformed into PQN4, an *E. coli* cell strain derived from LSR10 (MC4100, Δ*csgA*, λ(DE3), $Cam^R$) with the deletion of curli operon (ΔcsgBACEFG). Both pET21dCsgA-$\alpha$ and pET21dCsgA-$\gamma$ plasmids contained curli operon genes co-transcribed with the engineered csgA *i.e.*, *csgC*, *csgE*, *csgF* and *csgG*, which encodes the proteins necessary for biosynthesis of curli fibers. In both these plasmids, the *csgB* gene was deleted from the curli operon in order to secrete CsgA fused alpha or gamma proteins and self-assemble them to functional curli fibers (CsgA-$\alpha$ or CsgA-$\gamma$) in the culture medium, without anchoring (Δ*csgB*) to the bacterial surface.

**Plasmids used to make functional microbial ink:** The pET21dAzu plasmid was created similar to that of pET21dCsgA-$\alpha$ by replacing the gene of csgA-$\alpha$ with that of azurin, while we retained the SEC (N-terminal signal sequence) and N22 (N-terminal curli-specific targeting sequence) to allow secretion of azurin into the extracellular milieu. In case of pET21dCsgA-BPA plasmid (similar to pET21dCsgA-$\gamma$), the gene encoding BPA binding peptide was fused to the C-terminus of CsgA via a 36-amino-acid flexible linker. The MazF plasmid pL-MazF was derived from IPTG inducible plasmid pL6FO, and contains LacI repressor and kanamycin resistance genes, with a pLac promoter upstream of a *mazF* sequence derived from the genome of *E. coli* K-12 MG1655 (EcoCyc Accession EG11249).[1] All plasmids used for creating functional microbial inks were transformed into *E. coli* strain PQN4.

**Microbial production of engineered nanofibers:** pET21dCsgA-$\alpha$ and pET21dCsgA-$\gamma$ plasmids were transformed into PQN4 cells and streaked onto lysogeny broth (LB) agar plates containing 100 μg $ml^{-1}$ carbenicillin and 0.5% glucose (m $v^{-1}$) for catabolite repression of T7RNAP and incubated overnight at 37 °C. One colony was picked from each plate of PQN4CsgA-$\alpha$ and PQN4CsgA-$\gamma$, and cultured separately at 37 °C in 5 ml LB media, 100 μg $ml^{-1}$ carbenicillin and 2% glucose (m $v^{-1}$). The overnight cultures of PQN4CsgA-$\alpha$ and PQN4CsgA-$\gamma$ were transferred to a fresh 500 ml LB media containing 100 μg $ml^{-1}$ carbenicillin, and cultured upon IPTG induction either separately or together. This mono- or co-culture of PQN4CsgA-$\alpha$ and PQN4CsgA-$\gamma$ was placed in incubator shakers (225 rpm, 37 °C) for 48 h to express the engineered proteins CsgA-$\alpha$ and CsgA-$\gamma$, and self-assemble them into functional curli nanofibers.

**Preparation of CsgA-$\alpha$, CsgA-$\gamma$ and CsgA-$\alpha\gamma$ hydrogels (microbial ink):** The 48 h bacterial culture (500 ml) with CsgA-$\alpha$, CsgA-$\gamma$ or CsgA-$\alpha\gamma$ nanofibers was treated with 0.8 M (final concentration) guanidinium chloride (GdmCl) and stored at 4 °C for 1 h. Subsequently, the engineered nanofibers were concentrated on a polycarbonate membrane with 11 μm pores (EMD Millipore) using vacuum filtration. The engineered nanofibers deposited on filter membrane were treated with 8 M GdmCl for 5 min. After which, it was vacuum filtered and washed with sterile deionized (DI) water (50 ml) twice to remove the lysed bacterial debris. Then, the nucleic acids bound to the curli fibers were removed by 10 min incubation with nuclease solution (Benzonase, Sigma-Aldrich, 1.5 U $ml^{-1}$), which was followed by water washes (50 ml twice). Finally, the engineered curli nanofibers present on the filter membrane were incubated with 5% (m $v^{-1}$ in water) sodium dodecyl sulfate (gelator) for 5 min, followed by vacuum filtration and DI water washes (50 ml twice). The hydrogel of functional curli nanofibers (CsgA-$\alpha$, CsgA-$\gamma$ or CsgA-$\alpha\gamma$) formed on the filter membrane was scraped off and stored at 4 °C.

**Field-emission scanning electron microscopy (FESEM) sample preparation and imaging**: FESEM samples were prepared by fixing with 2% (w $v^{-1}$) glutaraldehyde and 2% (w $v^{-1}$) paraformaldehyde at room temperature, overnight. The samples were gently washed with water, and the solvent was gradually exchanged to ethanol with an increasing ethanol 15-minute incubation step gradient (25, 50, 75 and 100% (v $v^{-1}$) ethanol). The samples were then dried in a critical point dryer, placed onto SEM sample holders using silver adhesive (Electron Microscopy Sciences) and sputtered until they were coated in a 10-20 nm layer of Pt/Pd. Images were acquired using a Zeiss Ultra55 FESEM equipped with a field emission gun operating at 5-10 kV. Representative images from three independent samples were reported.

**Transmission electron microscopy (TEM) sample preparation and imaging**: 10 μl of the cultures of CsgA-$\alpha$, CsgA-$\gamma$ or CsgA-$\alpha\gamma$ was drop-casted onto formvar-carbon grids (Electron Microscopy Sciences), washed with DI water (thrice) and stained with 1% uranyl formate before analysis on a JEOL 1200 TEM. Representative images from three independent samples were reported.

**Optical Images**: Optical images were acquired using a Canon EOS Rebel SL3 Digital SLR Camera equipped with XIT 58 mm 0.43 Wide Angle Lens and XIT 58 mm 2.2x Telephoto Lens. Representative images from three independent samples were reported.

**Rheology studies of the hydrogels**: The viscoelastic properties of the hydrogels were determined using a Discovery Hybrid Rheometer-3 and TRIOS software (TA Instruments, New Castle, DE). Hydrogels were loaded between a Peltier plate and a 20 mm plate geometry. The excess sample was trimmed along the edge of the 20 mm plate. Samples were surrounded with mineral oil to prevent dehydration. Time sweep experiments were conducted under continuous oscillations at 0.1 Hz with an imposed shear strain of 0.5%. After the storage modulus (G') reached a plateau, a frequency sweep routine was applied under an oscillatory shear strain of 0.5% with the frequency increasing from 0.01 to 10 Hz. The storage modulus (G') and loss modulus (G") were recorded for both time and frequency sweep. The plateau storage modulus in the time sweep experiment was taken as the linear shear modulus. To probe the strain dependence of viscoelastic properties, samples were subjected to a strain sweep routine at a frequency of 0.1 Hz with the oscillatory strain increasing from 0.01 to 100%. To measure the viscosity of samples, samples were loaded between a Peltier plate and a 20 mm plate and a flow sweep routine was performed with shear rate increasing from 0.01 to 100 Hz. To determine the yield stress of samples, samples were subjected to a stress sweep routine at a frequency of 1 Hz with the oscillatory stress increasing from 1 to 200 Pa. The yield stress is defined as the oscillatory stress where the storage modulus decreases to 50% smaller than the linear shear modulus. Data obtained from at least three independent samples were reported.

**3D Printing of the hydrogels (microbial ink)**: Prior to printing, the bioinks were transferred into a 10-ml Luer-Lok™ syringe and centrifuged at 1,000 rpm for 2 min to remove any air bubbles. The standard needle used was a 27G tip with a premade ¼" blunt end from Fisnar. Bioprinting was performed using an ANET A8 (Shenzhen Anet Technology Co) 3D printer that was upgraded into an extrusion bioprinter. The bioprinting was first conducted at a range of feed rates (2-10 mm $s^{-1}$) and pressures (20-40 psi) to understand their effects on extrusion of the bioinks. For bioprinting of actual patterns, the feed rate was kept consistent at 2.5 mm $s^{-1}$, with a constant pressure of approximately 20 psi. The patterns were designed using Solidworks 3D design software (Dassault Systèmes SE). The 3D STL files were sliced using the slic3r engine from Repetier Host, which served as the main program for operating the extrusion bioprinter.[2]

**Print fidelity test for the hydrogels (microbial ink)**: Fidelity test namely filament collapse test was performed according to the previously published protocol.[3] A small structure with pillars featuring a series of spacing's (2L) was first 3D-printed. A single line (filament; green) of the bioink was extruded by the bioprinter from one side of the structure to the other end at a moderate nozzle moving speed of 5 mm $s^{-1}$, suspending the bioink between the pillars (purple) and bridging the gaps. Photographs were taken from the side of the structure post-bioprinting to measure the structural integrity under gravitational force ($F_g$), in the form of angles ($\theta$) of the overhung bioink fibers. Fidelity data fitting and theoretical modeling were conducted also in accordance with the previous report.[3] Data obtained from at least three independent samples were reported.

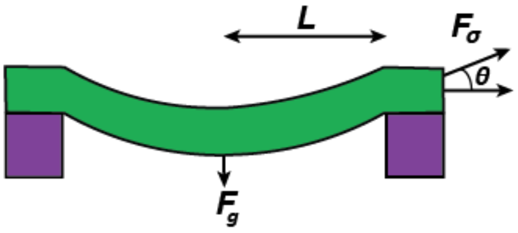

**Preparation of functional microbial ink:** The PQN4-Azu and PQN4-MazF cells were grown to OD 1 at 37 °C in 10 ml LB media, with 100 µg $ml^{-1}$ carbenicillin and 50 µg $ml^{-1}$ kanamycin, respectively. The 1 ml of microbial ink (CsgA-$\alpha\gamma$) was evenly spread on filter membrane and was incubated with 10 ml of bacterial culture (PQN4-Azu or PQN4-MazF) for 30 min to disperse the bacteria on the microbial ink and then vacuum filtered. In case of PQN4-BPA, the bacterial cells first were grown to OD 1 at 37 °C in 10 ml LB media with 100 µg $ml^{-1}$ carbenicillin and 2% glucose (m $v^{-1}$). Next, the cells were transferred to fresh LB media containing 100 µg $ml^{-1}$ carbenicillin and the CsgA-BPA fibers were expressed by induction with 0.1 mM IPTG. After 24 h of expression, the engineered PQN4-BPA biofilm was incorporated to 1 ml of microbial ink (CsgA-$\alpha\gamma$), as described above. The as-prepared microbial ink embedded with programmed bacterial cells/biofilms were utilized for 3D printing.

**Detection of secreted Azurin**: The 3D printed structures of microbial ink (CsgA-$\alpha\gamma$) embedded with PQN4-Azu cells were immersed in LB media containing 100 µg $ml^{-1}$ carbenicillin with or without 0.1 mM IPTG and incubated at 37 °C for 48 h. After 24 and 48 h of incubation, 0.5 ml of the culture media was collected for detection of azurin secreted by the microbial ink embedded with PQN4-Azu cells. The culture was concentrated on Amicon® Ultra-0.5 centrifugal filter (MiliporeSigma) to 20 µl. The protein present in concentrated supernatant were separated by NuPAGE 4–20% Tris-Glycine gels (ThermoFisher Scientific) and transferred to a nitrocellulose membrane using iBlot™ 2 Gel Transfer Device (ThermoFisher Scientific). The membrane was incubated with Goat Polyclonal Anti-Azurin Antibody (Origene) at 4 °C overnight. After washing, the membranes were incubated with a secondary antibody (rabbit anti-goat IgG-HRP (abcam) for 1 h at room temperature. Chemiluminescence was detected using a FluorChem M system (ProteinSimple). Data obtained from at least three independent samples were reported.

**BPA binding analysis**: The 3D printed structures of microbial ink (CsgA-$\alpha\gamma$) embedded with PQN4-BPA biofilm were immersed in water with or without 1 mM BPA and incubated for 24 h. After 12 and 24 h the water spiked with BPA was collected to test BPA binding by the microbial ink embedded with PQN4-BPA biofilm. The presence of BPA was detected using Agilent 6460 Triple Quad LC/MS and Agilent 1290 Infinity HPLC by Small Molecule Mass Spectrometry Facility at Harvard University, MA, USA. Data obtained from at least three independent samples were reported.

**Cell growth under MazF**: The 3D printed structures of microbial ink (CsgA-$\alpha\gamma$) embedded with PQN4-MazF cells were immersed with LB media consisting of 100 μg $ml^{-1}$ carbenicillin, with or without 0.1 mM IPTG at 37 °C for 6 h. The growth of PQN4-MazF cells in the microbial ink was monitored by collecting ~200 mg of the ink after 0, 2, 4 and 6 h. The collected microbial inks were transferred to the Eppendorf tubes and 1 ml of LB media was added to resuspend the ink, which was serially diluted, and plated on carbenicillin selective plates to obtain their colony forming unit (CFU) counts. Data obtained from at least three independent samples were reported.

**Statistics and reproducibility**: All experiments presented in this Article were repeated at least three times (n ≥ 3) on distinct samples, as clearly specified in the figure legends or the relevant Methods sections. In all cases, data are presented as the mean and standard deviation. GraphPad Prism 8 software was used for plotting and analyzing data. For micrographs and optical images, we present representative images.

### References (Methods only)


1 Kan, A., Birnbaum, D. P., Praveschotinunt, P. & Joshi, N. S. Congo Red Fluorescence for Rapid In Situ Characterization of Synthetic Curli Systems. *Appl Environ Microbiol* **85**, doi:10.1128/AEM.00434-19 (2019).

2 Ying, G. L. *et al.* Aqueous Two-Phase Emulsion Bioink-Enabled 3D Bioprinting of Porous Hydrogels. *Adv Mater* **30**, e1805460, doi:10.1002/adma.201805460 (2018).

3 Ribeiro, A. *et al.* Assessing bioink shape fidelity to aid material development in 3D bioprinting. *Biofabrication* **10**, 014102, doi:10.1088/1758-5090/aa90e2 (2017).

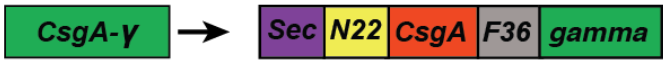


| Name | Amino acid sequence |
| --- | --- |
| Sec | MKLLKVAAIAAIVFSGSALA |
| N22 | GVVPQYGGGGNHGGGGNNSGPN |
| CsgA | GVVPQYGGGGNHGGGGNNSGPNSELNIYQYGGGNSALALQTDAR NSDLTITQHGGGNGADVGQGSDDSSIDLTQRGFGNSATLDQWNGK NSEMTVKQFGGGNGAAVDQTASNSSVNVTQVGFGNNATAHQY |
| Linker F36 | GGSGSSGSGGSGGGSGSSGSGGSGGGSGSSGSGGSG |
| gamma | DAGDAFDGFDFGDDPSDKFFTSHNGMQFSTWDNDNDKFEGNCAE QDGSGWWMNKCHAGHLNGVYYQGGTYSKASTPNGYDNGIIWAT WKTRWYSMKKTTMKIIPFNRLTIGEGQQHHLGGAKQAGDV |

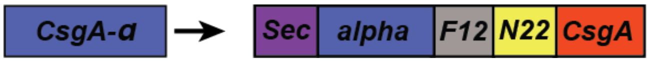


| Name | Amino acid sequence |
| --- | --- |
| Sec | MKLLKVAAIAAIVFSGSALA |
| alpha | GPRVVERHQSA |
| N22 | GVVPQYGGGGNHGGGGNNSGPN |
| Linker F12 | GSGGSGGSGGSG |
| CsgA | GVVPQYGGGGNHGGGGNNSGPNSELNIYQYGGGNSALALQTDAR NSDLTITQHGGGNGADVGQGSDDSSIDLTQRGFGNSATLDQWNGK NSEMTVKQFGGGNGAAVDQTASNSSVNVTQVGFGNNATAHQY |

**Supplementary Table 1. Amino acid sequences of engineered curli nanofibers of CsgA-$\alpha$ and CsgA-$\gamma$ that produce the microbial ink CsgA-$\alpha\gamma$.**

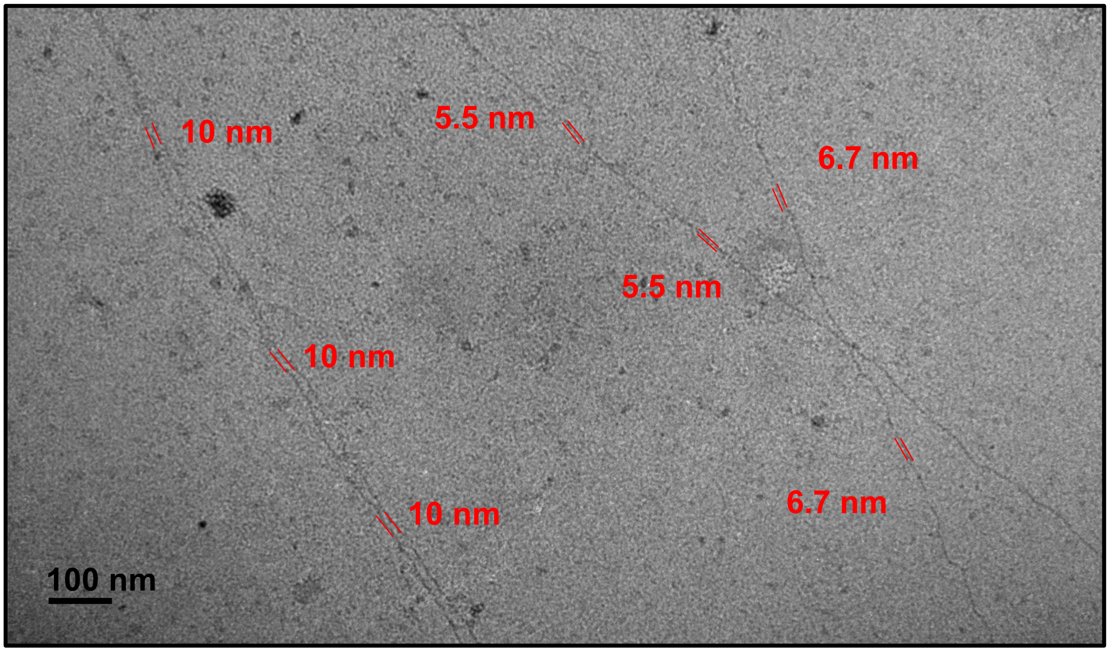


**Supplementary Figure 1. TEM image of a culture of CsgA-$\alpha\gamma$, showing the diameters of the nanofibers.** The 10 nm fiber could be attributed to CsgA-$\alpha\gamma$ formed by the supramolecular crosslinking of alpha (knob) and gamma (hole) modules of CsgA-$\alpha$ and CsgA-$\gamma$, respectively. The nanofibers with 5.5 and 6.7 nm corresponds to CsgA-$\alpha$ and CsgA-$\gamma$, respectively.

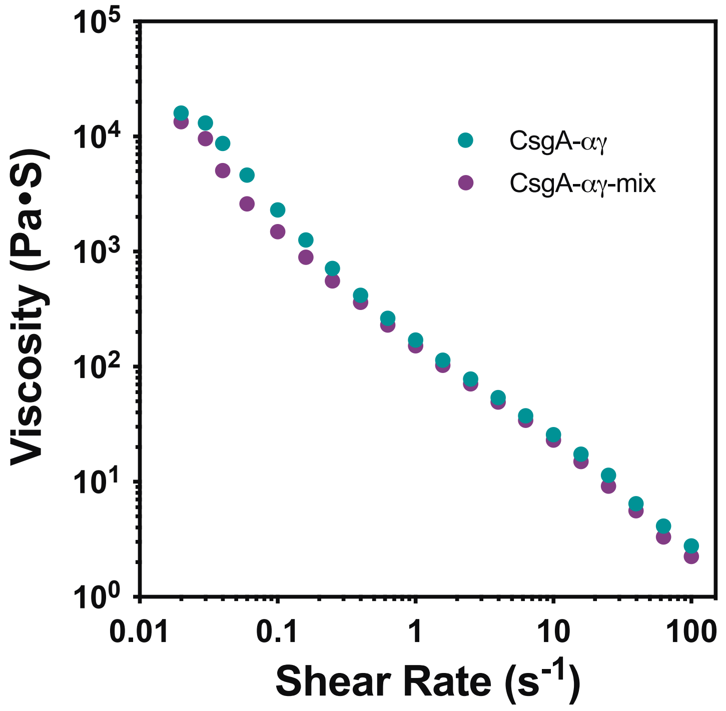


**Supplementary Figure 2. Shear-thinning behavior of CsgA-$\alpha\gamma$ and CsgA-$\alpha\gamma$-mix.** The decreasing viscosity with increasing shear rate indicates the shear-thinning property of CsgA-$\alpha\gamma$ (co-culture of CsgA-$\alpha$ and CsgA-$\gamma$) and CsgA-$\alpha\gamma$-mix (1 h mixing of separately cultured CsgA-$\alpha$ and CsgA-$\gamma$). The viscosity of CsgA-$\alpha\gamma$-mix is similar to that of CsgA-$\alpha\gamma$.

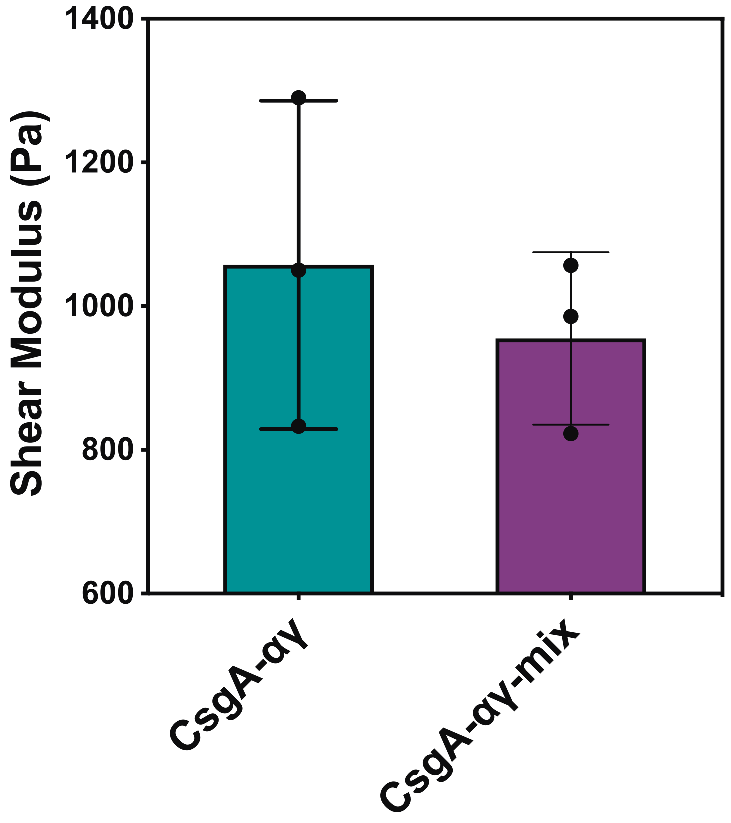


**Supplementary Figure 3. Shear modulus of of CsgA-$\alpha\gamma$ and CsgA-$\alpha\gamma$-mix.** The shear modulus of CsgA-$\alpha\gamma$-mix (1 h mixing of separately cultured CsgA-$\alpha$ and CsgA-$\gamma$) is similar to that of CsgA-$\alpha\gamma$ (co-culture of CsgA-$\alpha$ and CsgA-$\gamma$).

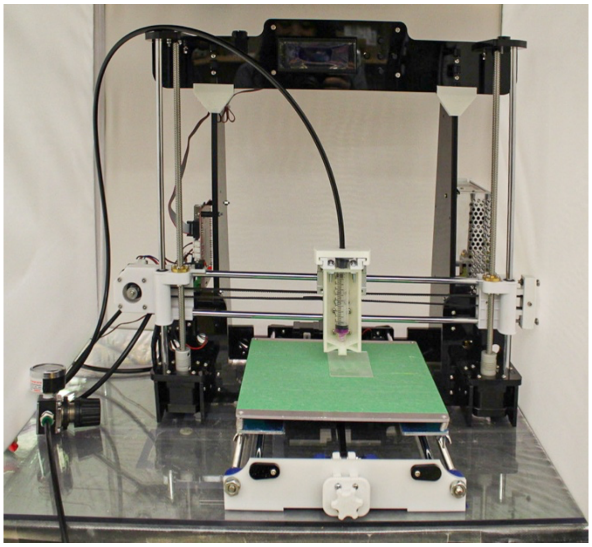

**Supplementary Figure 4. Optical image of the customized 3D printer.**

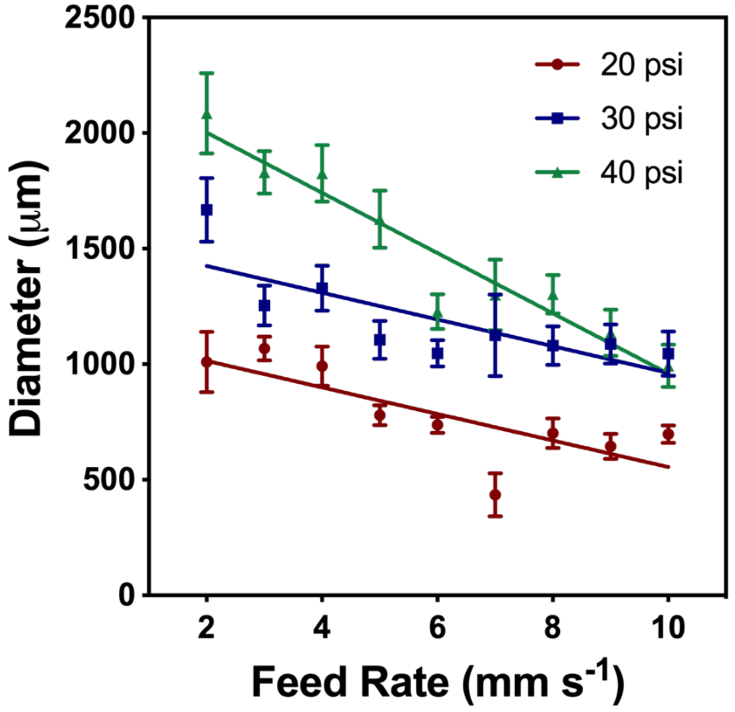


**Supplementary Figure 5. Printing performance of CsgA-$\alpha$.** Plot shows the line width of CsgA-$\alpha$ hydrogel-based bioink at various feed rates (2-10 mm $s^{-1}$) and pressures (20-40 psi).

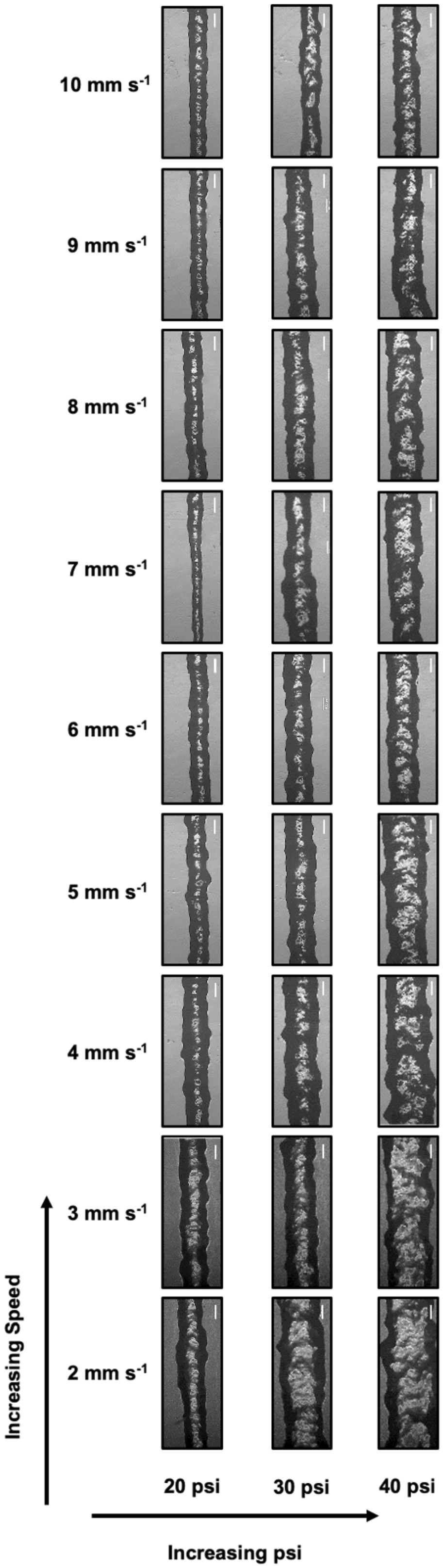


**Supplementary Figure 6. Printing performance of CsgA-$\alpha$.** Optical images show the line width of CsgA-$\alpha$ hydrogel-based bioink at various feed rates (2-10 mm $s^{-1}$) and pressures (20-40 psi). Scale bar 500 $\mu$m.

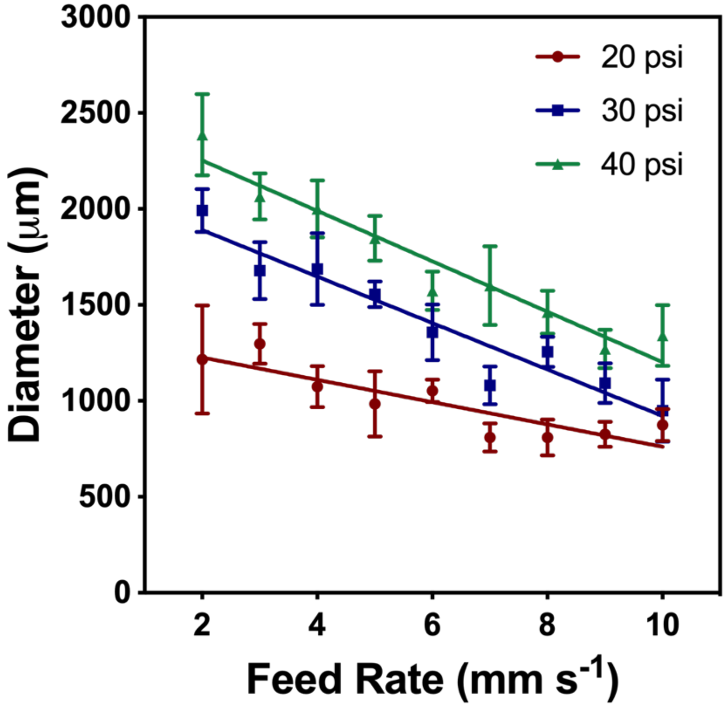


**Supplementary Figure 7. Printing performance of CsgA-$\gamma$.** Plot shows the line width of CsgA-$\gamma$ hydrogel-based bioink at various feed rates (2-10 mm $s^{-1}$) and pressures (20-40 psi).

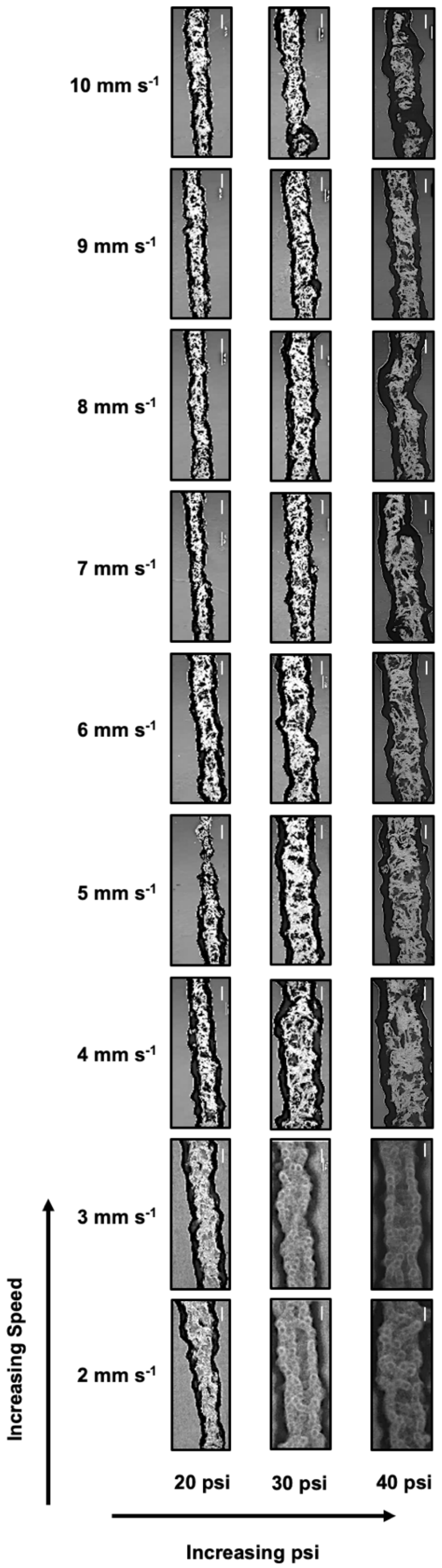


**Supplementary Figure 8. Printing performance of CsgA-$\gamma$.** Optical images show the line width of CsgA-$\gamma$ hydrogel-based bioink at various feed rates (2-10 mm $s^{-1}$) and pressures (20-40 psi). Scale bar 500 $\mu$m.

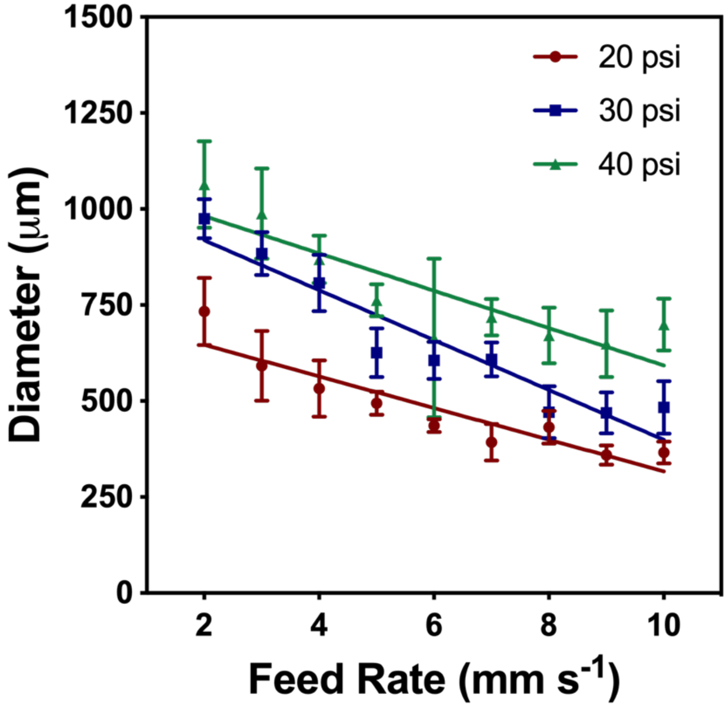


**Supplementary Figure 9. Printing performance of CsgA-$\alpha\gamma$.** Plot shows the line width of CsgA-$\alpha\gamma$ hydrogel-based bioink at various feed rates (2-10 mm $s^{-1}$) and pressures (20-40 psi).

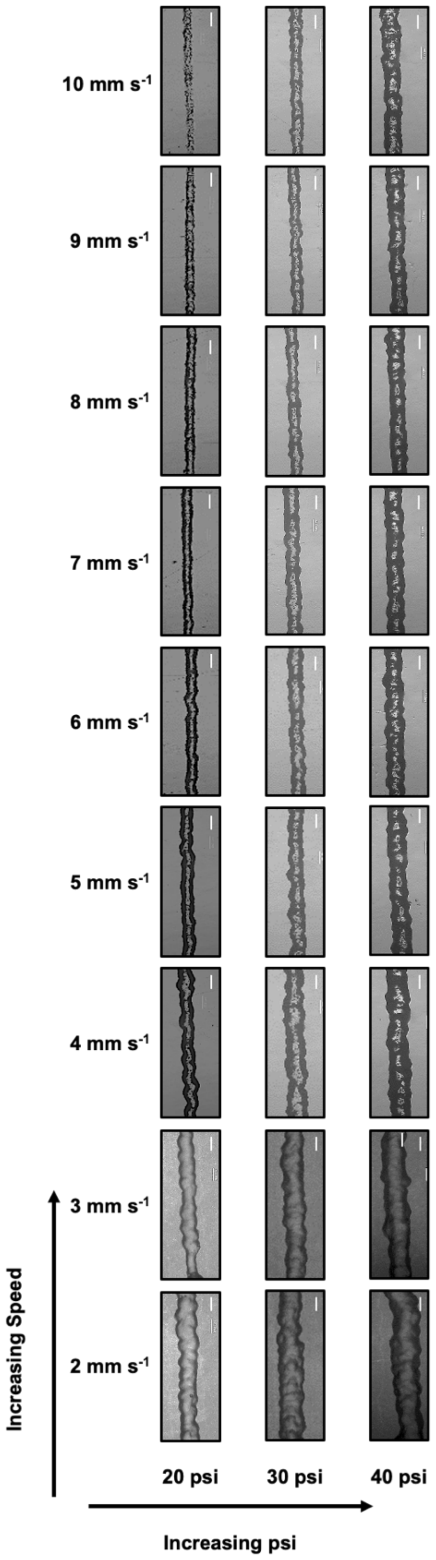


**Supplementary Figure 10. Printing performance of microbial ink CsgA-$\alpha\gamma$.** Optical images show the line width of CsgA-$\alpha\gamma$ bioink at various feed rates (2-10 mm $s^{-1}$) and pressures (20-40 psi). Scale bar 500 $\mu$m.

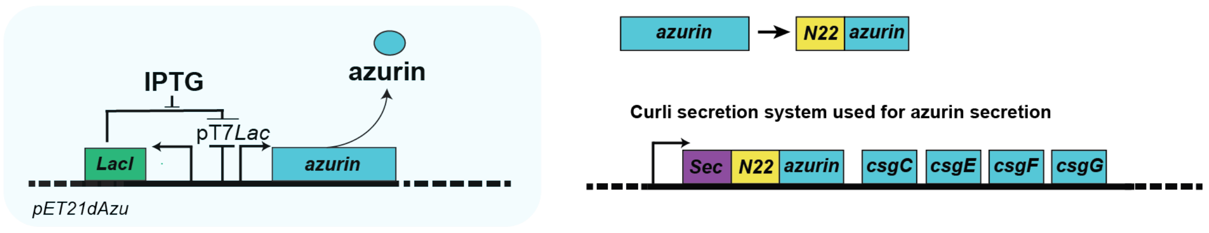


| Name | Amino acid sequence |
|---|---|
| **Sec** | MKLLKVAAIAAIVFSGSALA |
| **N22** | GVVPQYGGGGNHGGGGNNSGPN |
| **Azurin**[1] | AECSVDIQGNDQMQFNTNAITVDKSCKQFTVNLSHPGNLPKNVMG HNWVLSTAADMQGVVTDGMASGLDKDYLKPDDSRVIAHTKLIGSG EKDSVTFDVSKLKEGEQYMSFCTFPGHSALMKGTLTLK |

**Supplementary Figure 11. Genetic design of pET21dAzu to secrete azurin.** The anticancer biologic drug azurin was secreted via curli secretion system in PQN4-Azu cells by using pET21dAzu plasmid. The table shows the amino acid sequence of azurin.

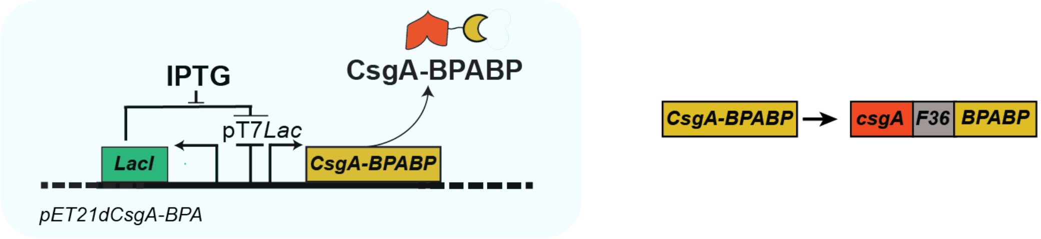


| Name | Amino acid Sequence |
|---|---|
| **CsgA** | GVVPQYGGGGNHGGGGNNSGPNSELNIYQYGGGNSALALQTDAR NSDLTITQHGGGNGADVGQGSDDSSIDLTQRGFGNSATLDQWNGK NSEMTVKQFGGGNGAAVDQTASNSSVNVTQVGFGNNATAHQY |
| **Linker F36** | GGSGSSGSGGSGGGSGSSGSGGSGGGSGSSGSGGSG |
| **BPABP**[2] | KSLENSY |

**Supplementary Figure 12. Genetic design of pET21dCsgA-BPA to bind BPA.** BPA binding peptide (BPABP) was genetically grafted to the CsgA protein to obtain PQN4-BPA biofilm. The table shows the amino acid sequence of CsgA-BPABP.

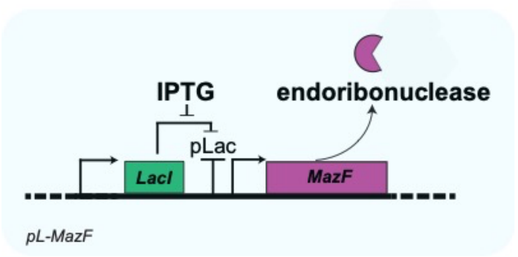


| Name | Amino acid sequence |
|---|---|
| MazF | MVSRYVPDMGDLIWVDFDPTKGSEQAGHRPAVVLSPFMYNNKTG MCLCVPCTTQSKGYPFEVVLSGQERDGVALADQVKSIAWRARGAT KKGTVAPEELQLIKAKINVLIG |

**Supplementary Figure 13. Genetic design of pL-MazF to secrete MazF.** The toxin MazF was expressed in PQN4-MazF cells by using pL-MazF plasmid. The table shows the amino acid sequence of MazF.
